# Memory schema reorganization induced by the deliberate processing in the executive network supported by widespread amplified activity

**DOI:** 10.64898/2025.12.30.697002

**Authors:** Hiroki Kurashige, Jun Kaneko, Kenji Matsumoto

## Abstract

Schema accommodation is the reorganization of a preexisting memory schema to deeply accept new information incongruent with it. Although it is crucial for flexible intelligence, it occurs infrequently, making its neural basis difficult to study. To overcome this, we conducted an fMRI experiment using a newly developed experimental paradigm, the reversal description task. The results suggest that the schema changes that occurred during the task were global reorganizations rather than local adjustments. Moreover, the executive network responsible for deliberate processing played a central role, with the support of widespread amplified activity. Furthermore, reinforcement learning-based control implemented in these neural substrates, in cooperation with the caudate nucleus, emerged as a candidate mechanism for the deliberative schema update. These findings indicate that memory reorganization involves not only automatic but also deliberate processes, in which the desired global structure is explored through the control operation performed by these neural bases.

**Teaser:** Memory reorganization is performed through deliberation-related neural computations supported by brain-wide amplified activity.

## Introduction

We can think about something because we have knowledge and memories of it. This is neither an isolated datum nor a collection of scattered data. Because memory functions as a complex but integrated system, it should be structured systematically. Such a structure is called memory schema, which is an organized framework of knowledge/memory formed in the brain through experiences, including online and offline processes of learning and thought (*1–4*). Its detailed definition differs slightly depending on the discipline; however, for the purposes of this paper it is sufficient to consider it as a network of various things interconnected by various relationships. The structure of a schema affects recognition and thoughts, thereby determining behavior (*5–8*). In other words, we can behave only as our own schema structures allow. Therefore, to understand what kind of computational beings humans are, we need to know the organizing principles governing the formation and updating of structures through experience.

Generally, the acquisition of new information depends on preexisting memory schemata (*9–14*). Because this provides the organization of the next schema structure, it is a self-organized recursive process. This schema-dependent acquisition of novel knowledge is of two types: schema assimilation and schema accommodation (Fig. 1A) (*1*, *2*, *15*). Schema assimilation is a phenomenon in which new information is assimilated without substantial changes to the preexisting schema itself. A characteristic of this phenomenon is that the acquisition of schema-congruent information is facilitated, as is often experienced in our daily lives. This makes schema assimilation a more frequently occurring process. Phenomena that occur frequently and automatically in natural situations can also be easily observed in the laboratory. Indeed, the properties of schema assimilation are steadily being elucidated. In particular, the underlying neural mechanisms are becoming better understood. The central role of the ventromedial prefrontal cortex (vmPFC) is almost established (*3*, *4*, *9*).

**Fig. 1.**
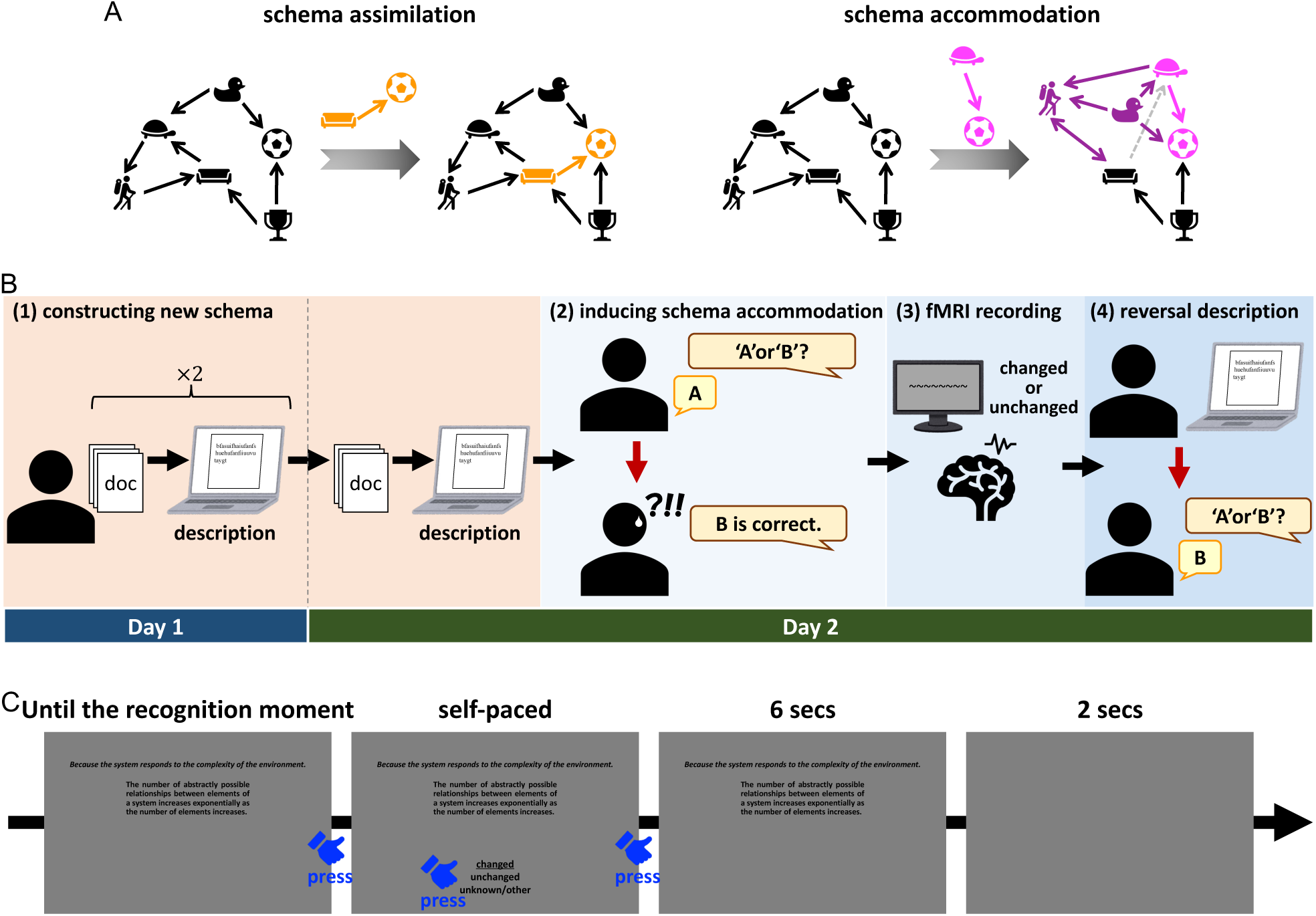
Background and experimental setting. **(A)** Schema assimilation and accommodation. In assimilation, new information congruent with the existing schema (orange) is incorporated without substantial changes to the schema itself. In accommodation, substantial reorganization of the schema (dark magenta) is required to incorporate information incongruent with the schema (light magenta). **(B)** Reversal description task. (1) Participants read the text while taking notes, then described their understanding, referring only their notes (without access to the original document). (2) They answered a two-choice question based on their interpretation. They were then informed that the opposite answer was correct and were instructed, during the fMRI task, to revise their interpretation so that the opposite response would become their own; they were also instructed that, after the session, they would describe their understanding again. (3) During the fMRI session, sentences constituting the document were presented. Participants rethought each sentence and reported whether its interpretation had changed. (4) After the recording, they described their understanding of the document again and then answered the same two-choice question again. **(C)** fMRI task procedure. Participants pressed a button at the moment they internally recognized whether the interpretation of each sentence had changed or not. The screen then changed, and they reported their decision, followed by a period of passive viewing of the sentence and a blank period. The blue finger icons were not shown to participants and the texts were presented in Japanese.

In contrast, the neural mechanisms underlying schema accommodation remain largely unknown. When one encounters information that is incongruent with one’s existing schema, its structure may need to be reorganized to incorporate the information into it. Schema accommodation is such a reorganization phenomenon on a memory schema. This is vital for flexible, adaptive, and creative memory. However, the requirement for such a substantial structural change in the schema makes it resistant, resulting in this being less likely to occur, as many psychological studies have shown (*16–18*). This has made it difficult to elucidate the neural mechanisms that induce and/or support schema accommodation.

If there is no substantial reorganization of an existing schema, it is not schema accommodation. For example, constructing a new memory is not a schema accommodation (it is, at best, just the formation of a new schema), nor is it simply adding new information to an existing memory schema (i.e., schema assimilation). Furthermore, even when an update to an existing schema occurs, it should not be schema accommodation if it is only a localized change, such as the modification of a single association. Schema accommodation should involve changes in the global structure of memory (i.e., reorganization).

Neurophysiological findings regarding schema and memory updating are progressively accumulating (*19–21*). In most studies along this line, participants were asked to first memorize artificial relations designed by experimenters, such as paired associations, and then modify parts of their memories (*22–24*). One experiment used a task with complex room images, in which multiple objects were placed as targets to be learned (*25*). However, even in this case, the position of each object was arbitrary. This means that, as in most other studies, this updating of memory was just the sum of the local modifications of individualized items that were unrelated to each other. Therefore, these memory schema updates do not necessarily meet the aforementioned conditions for schema accommodation; therefore, it cannot be said that these studies investigate neural mechanisms of schema accommodation.

In addition, several studies have investigated the neural phenomena involved in updating memory schemata under more naturalistic conditions (*26–29*). We expect the items in these schemata to be interrelated, similar to those in everyday life. Thus, updating memories examined in such experiments may meet the conditions for schema accommodation. Using a real movie, one study showed that the neural representation of a scene changes after the presentation of a twist (*27*). However, this is, at best, a report of a phenomenon that occurred as a result of schema accommodation and does not examine its mechanism. Experiments were also conducted in which, first, sets of seemingly unrelated scenes were presented using virtual reality, and then, events that were linked between them were presented (*26*, *28*, *29*). One such experiment showed that perturbing activity in the left angular gyrus with transcranial magnetic stimulation inhibited linking (*29*). However, what was examined in that study was at most the linking between two individualized schemata, and it remains unclear whether there was any within-schema reorganization (i.e., schema accommodation) occurred in either schema. Some relationships between these experiments and the present study are revisited in the Discussion section.

Therefore, in the present study, we aim (i) to elucidate the neural mechanisms that either induce or support schema accommodation (and in some cases both), and (ii) to characterize the types of psychological and computational processes involved in the mechanisms, and (iii) to provide an insight into fundamental understanding of the dynamical laws that govern schema structures and their temporal evolution. To this end, we conducted a functional magnetic resonance imaging (fMRI) experiment using a newly developed task paradigm, the reversal description task (Fig. 1B), which effectively induces schema accommodation and enables the investigation of its neural substrates.

## Results

To investigate the neural mechanisms that induce and/or support schema accommodation, we conducted an fMRI experiment using the newly developed reversal-description task (Fig. 1B). In this task, each participant was asked to read a document that was so esoteric that it required active interpretation to make sense of it and then write a text summarizing his/her understanding. This reading and writing set was repeated three times: twice on the first day and once on the second day. The document was extracted from an actual theoretical sociology book (*30*) that was pedantic but insightful. This procedure forces them to construct a new schema in their minds that is sufficiently rich and robust. Then, the participants answered a two-choice question (“A” vs. “B”). Their answers should have differed according to the interpretation of the document (i.e., their schemata). After this, they were told that the opposite of each answer was “correct.” They were also instructed that they needed to change their interpretation so that the “correct answer” would be their own answer through the task during the MRI measurement. In addition, the participants were informed that they would write a text summarizing their understanding of the document based on the changed interpretations after the measurement. During the fMRI session, sentences extracted from the document were presented to the participants (Fig. 1C). Each participant read and rethought each sentence and reported whether the interpretation of the sentence (rather than the interpretation of the entire document or the answer to the question above) had changed. Trials that were reported to have changed were considered to have undergone some changes in the schema. After the MRI session, the participants rewrote their current understanding of the document and then answered the same two-choice question as before. See the Materials and Methods for more details and reasoning regarding the procedure.

### Behavioral results suggest global changes in the schema structure in the accommodation group

As explained above, schema accommodation reorganizes a schema to accept incongruent information. In our analysis, this is defined operationally. That is, participants whose answers flipped before and after the fMRI measurement (i.e., “A”→“B” or “B”→“A”) were the accommodation group, and those who did not flip (i.e., “A”→“A” or “B”→“B”) were the no-accommodation group. However, the resultant classification using this definition also successfully captures the mechanistic aspects of schema accommodation. This is demonstrated in the subsequent analyses.

Based on this definition, 22 of the 34 participants were classified into the accommodation group and 11 into the no-accommodation group. Since participants who gave a confidence level of “1. Just guessing” for at least one of the answers could not be classified into either group, they were excluded from further analyses. One participant was excluded on the basis of this criterion. Thus, the results showed that 2/3 of the participants analyzed underwent schema accommodation, while 1/3 did not. This is reasonable, given schema accommodation’s resistance to induction, while still yielding enough instances to permit investigation of its neural mechanisms.

Hereafter, we refer to trials in which participants responded that their interpretation of the presented sentences “changed” during the fMRI task as interpretation-changed trials, and trials in which they responded “unchanged” as interpretation-unchanged trials. Our primary interest here was to characterize the differences in brain activity during these two types of trials and their psychological and computational implications. We compared the time taken to decide whether or not the interpretation of a sentence had changed between the trial types. In the accommodation group, the mean duration of the interpretation-changed trials was significantly longer than that in the interpretation-unchanged trials, but there was no significant difference in the no-accommodation group (Figs. 2A and B; Wilcoxon signed-rank test, *n* = 22, *T* = 12, *p* = 3.34 × 10^−5^ for the accommodation group, *n* = 11, *T* = 16, *p* = 0.147 for the no-accommodation group). This is due to the fact that the mean decision duration of the interpretation-unchanged trials in the accommodation group was shorter than that of the others. Indeed, unpaired comparisons indicated that within the accommodation group, the mean durations of interpretation-changed trials were longer than those of interpretation-unchanged trials (Fig. 2C; Mann-Whitney U test, *n*_1_ = *n*_2_ = 22, *U* = 359, *p* = 0.00625), whereas neither the mean duration of interpretation-changed trials in the no-accommodation group (*n*_1_ = 22, *n*_2_ = 11, *U* = 106, *p* = 0.580) nor that of interpretation-unchanged trials in the no-accommodation group (*n*_1_ = 22, *n*_2_ = 11, *U* = 130, *p* = 0.745) differed from the mean duration of the interpretation-changed trials in the accommodation group. These results suggest that, at least in the accommodation group, there were some psychological and/or computational differences between information processing in the two trial types and that the results of subsequent imaging analyses for the group may reflect these differences. As the decision duration of interpretation-unchanged trials in the accommodation group was shorter than that in the others, one might consider that participants cut corners here more than in the other cases. However, we note that this is probably not the case, as later fMRI analyses (in particular, the analysis of beta estimates) suggest.

**Fig. 2.**
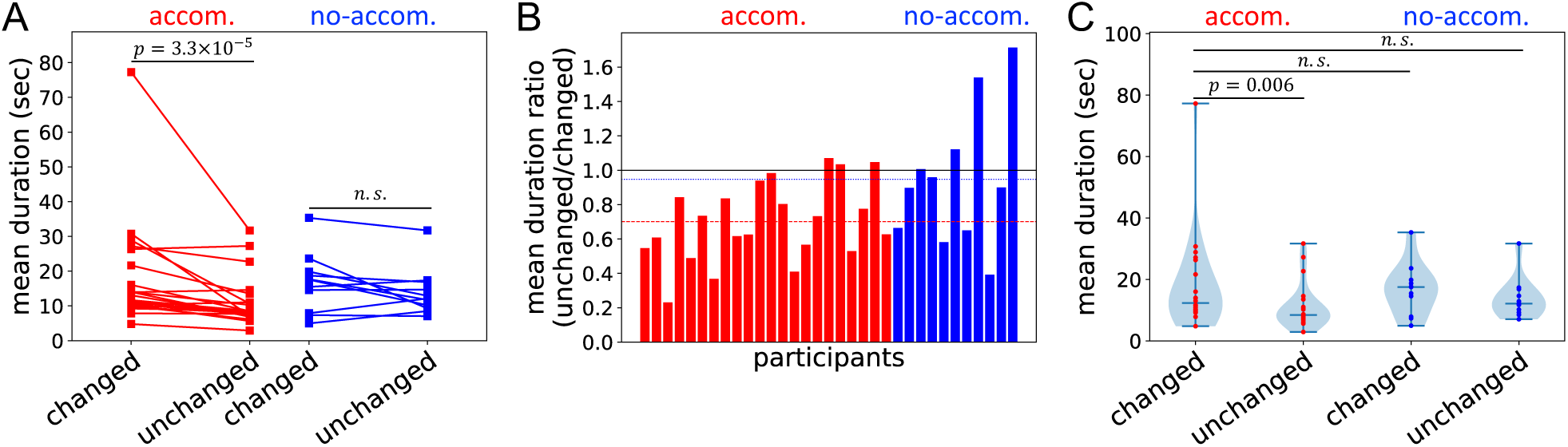
Behavior comparison between the accommodation and no-accommodation groups. **(A)** Paired comparisons of durations until participants recognized whether the interpretation of a presented sentence had changed. In the accommodation group, the mean duration for interpretation-changed trials was significantly longer than that for interpretation-unchanged trials (red). No such difference was observed in the no-accommodation group (blue). Each line represents an individual participant. The vertical axis indicates the mean duration per participant. **(B)** Ratio of durations for interpretation-unchanged trials to interpretation-changed trials. The mean ratio was approximately 0.701 for the accommodation group (red dashed line) and approximately 0.948 for the no-accommodation group (blue dotted line). Each bar represents an individual participant. **(C)** Unpaired comparisons of the mean duration of interpretation-changed trials in the accommodation group (leftmost) with those in other conditions. The mean duration of interpretation-unchanged trials in the accommodation group was significantly shorter than that of interpretation-changed trials in the same group. No significant differences were observed when the mean duration of interpretation-changed trials in the accommodation group was compared with those of interpretation-changed and interpretation-unchanged trials in the no-accommodation group. Each dot represents an individual participant, and the vertical axis indicates the mean duration per participant.

Next, the differences between the sentences reported as “changed” in interpretation and those reported as “unchanged” were analyzed using natural language processing and network theory methods. First, we measured the semantic similarity between the 40 sentences presented during the fMRI task using the BERTScore (*31*), constructed a network with similarity as edge weights, and obtained the eigenvector centrality of each sentence (Figs. 3A and B). In both the accommodation and no-accommodation groups, the sentences reported as “changed” had higher mean centrality than those reported as “unchanged” (Wilcoxon signed-rank test, *n* = 22, *T* = 14, *p* = 5.25 × 10^−5^ for the accommodation group, *n* = 11, *T* = 5, *p* = 0.00977 for the no-accommodation group). However, this did not differ between the groups (mixed ANOVA, *F*(1,31) = 6.19 × 10^−4^, *p* = 0.980). In addition, we examined whether interpretation-changed and interpretation-unchanged sentences differed in semantic similarity to the two sentences used in the question (which had approximately equal semantic centrality; Fig. 3C). We found that the former had a higher similarity to the question sentences than the latter (Fig. 3D; Wilcoxon signed-rank test, *n* = 22, *T* = 7, *p* = 9.06 × 10^−6^ for the accommodation group; *n* = 11, *T* = 7, *p* = 0.0186 for the no-accommodation group). However, in this case as well, there were no significant differences between the groups (mixed ANOVA, *F*(1,31) = 0.103, *p* = 0.751). Thus, since no difference exists between the accommodation and no-accommodation groups in terms of localized item-wise change, it is suggested that our schema accommodation should be due to actual global changes in the schema structure, even though we defined it operationally. Therefore, the differences in brain activity between the two groups presented in the following sections are considered to show the neural mechanisms that induce and/or support global changes in schema structure. See the Discussion section for a fuller explanation of this conclusion, where these behavioral results are considered together with the subsequent fMRI findings to provide stronger support for it.

**Fig. 3.**
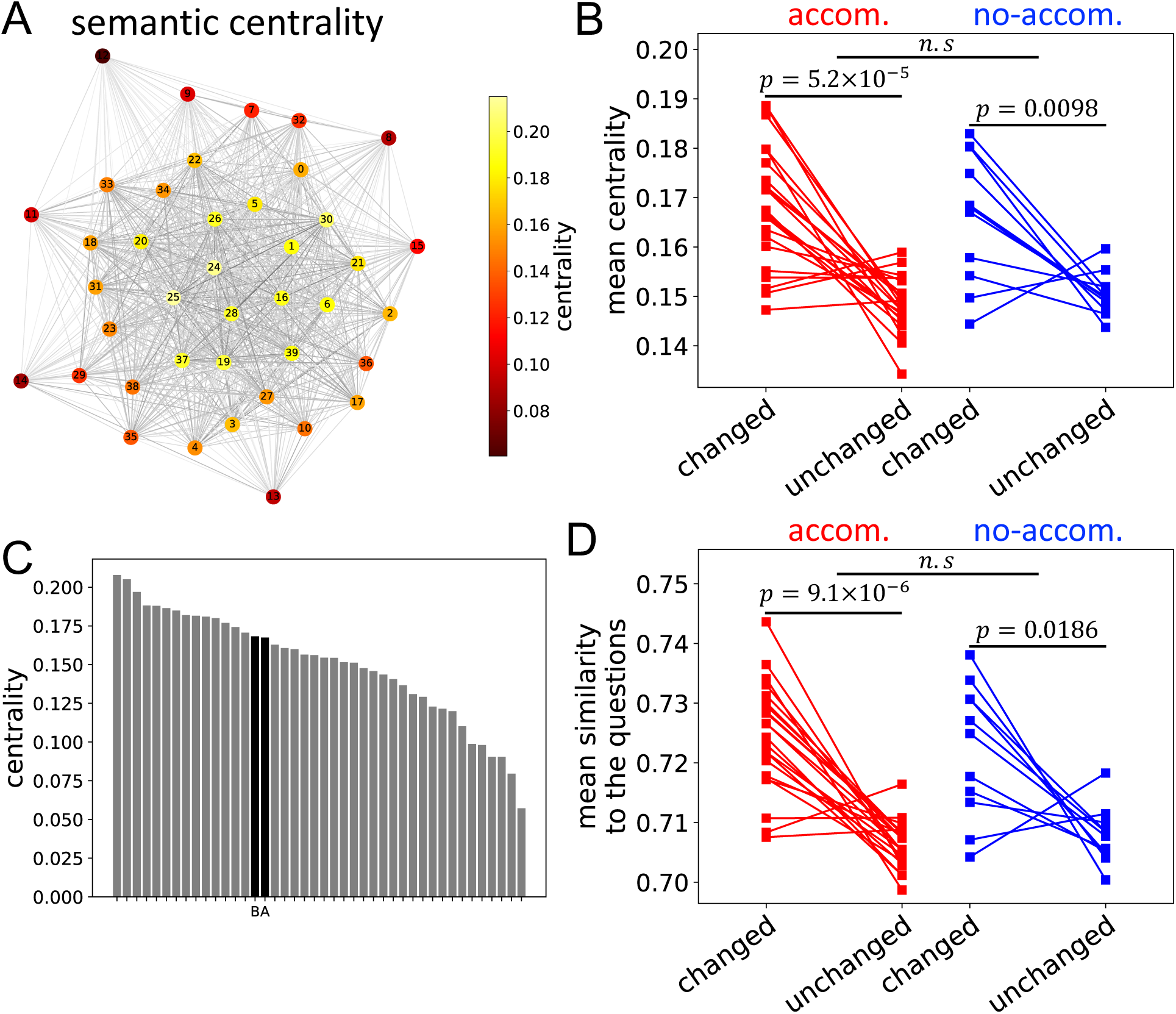
Analysis of semantic associations between sentences presented in an fMRI session. **(A)** A semantic network of the presented sentences, with edge weights derived from BERTScore similarity scores. Node coloring reflects eigenvector centrality values, and numbers indicate the order of sentence appearance in the document. **(B)** Comparisons of semantic centrality between sentences presented in interpretation-changed and interpretation-unchanged trials for the accommodation and no-accommodation groups. In both groups, sentences from interpretation-changed trials were significantly more central. This pattern did not differ between groups. The vertical axis represents the mean centrality per participant. **(C)** Semantic centrality of sentences presented during the fMRI session and those used in the two-choice questions. The same centrality analysis was extended to include the question sentences. Sentences are sorted in descending order of centrality. Each bar represents a sentence, with “A” and “B” indicating the question sentences. **(D)** Comparisons of sentence similarity to the two-choice question sentences between interpretation-changed and interpretation-unchanged trials. In both groups, sentences from interpretation-changed trials were significantly more similar to the question sentences. This pattern did not differ between groups. The vertical axis represents the mean similarity per participant.

### Schema accommodation can be induced by deliberate processing implemented in the executive network

To explore the neural mechanisms that induce and/or support schema accommodation, we compared fMRI signals during interpretation-changed and interpretation-unchanged trials. From the whole-brain analysis in the accommodation group, we found four voxel clusters that showed significantly stronger signals during interpretation-changed trials (Fig. 4 and Table S1): the left middle frontal gyrus (MFG), left inferior parietal lobule (IPL) including the supramarginal and angular gyri (roughly corresponding to its PFm and Pga subregions, respectively), left medial part of the superior frontal gyrus (medial SFG), and left lateral part of the superior frontal gyrus (lateral SFG). In a certain taxonomy, voxel clusters in the left MFG and left IPL constitute the left frontoparietal (FP) network (*32*, *33*). The voxel cluster in the left medial SFG can be considered a part of the left cingulo-opercular (CO) network (*34*); however, according to a well-known cortical atlas (*35*), it is mostly included in the left FP network. The FP and the CO networks constitute the executive network (*36–38*). Note that, in many cases, “executive function,” “central executive function,” “executive control,” and “cognitive control” are synonymous (see Ref. (*38*)). In addition, the executive, central executive, cognitive control, and multiple-demand networks are treated as synonyms (see Ref. (*39*)). Therefore, in this study, we follow these usages and do not distinguish between them. The executive network involves intelligent higher-order information processing (i.e., deliberation), such as mathematical reasoning and planning (*40–42*). This suggests that schema accommodation is inducible by deliberate processing since greater activity in the network of the accommodation group was observed during interpretation-changed trials.

**Fig. 4.**
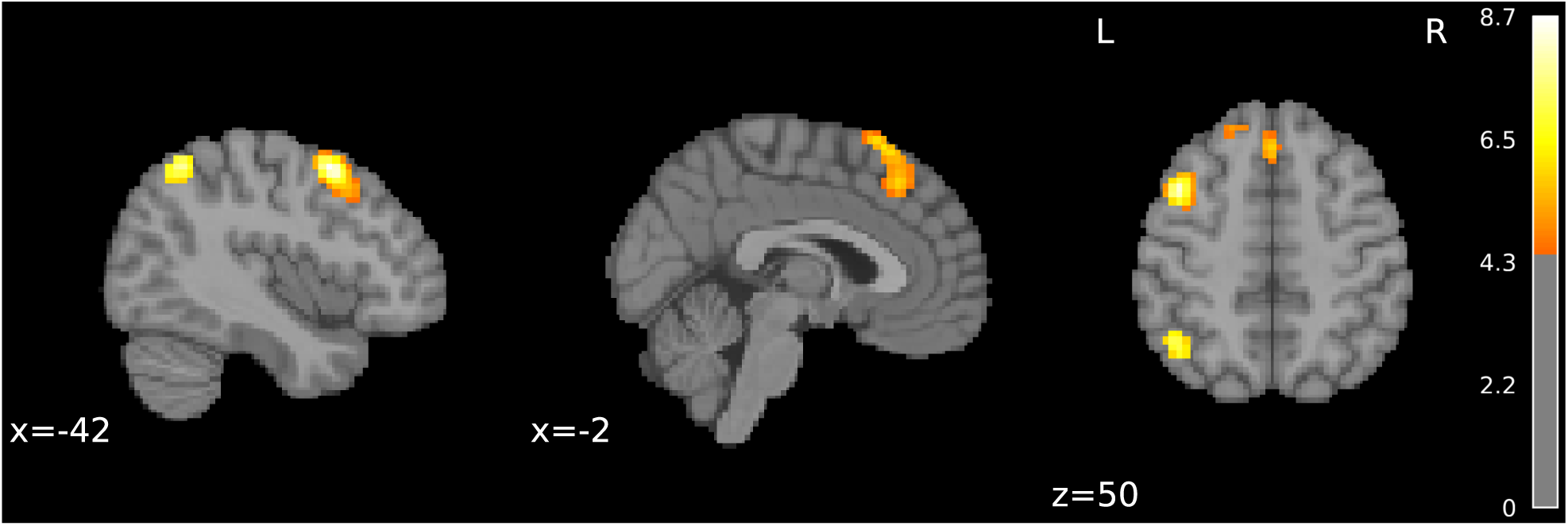
The executive network is involved in the reorganization of the schema in the accommodation group. Brain regions showing significantly greater activity during interpretation-changed trials compared to interpretation-unchanged trials in the accommodation group are displayed ( *p* < 0.05, family-wise error corrected). These regions were primarily located within the executive network, including the left middle frontal gyrus, the left inferior parietal lobule, the medial part of the left superior frontal gyrus—comprising the left frontoparietal network—and the lateral part of the left superior frontal gyrus. Voxel-wise t-values are represented using a color-coded scale.

In contrast, the no-accommodation group showed no clusters in which the fMRI signals were significantly greater in interpretation-changed trials than in interpretation-unchanged trials. However, owing to the difference in the number of participants between the two groups, the statistical test for the no-accommodation group had lower power. Therefore, we conducted the same test again for the no-accommodation group using the clusters identified in the previous analysis of the accommodation group as a mask (i.e., doing the so-called small-volume correction). Even in this case, no regions were significantly more strongly activated in the interpretation-changed trials, except for a single slightly significant voxel (*p* = 0.0493, *x* = 0, *y* = 23, *z* = 50). Thus, when schema accommodation did not occur, the increase in activity in the executive network was negligible (if any). In addition, as described in the previous paragraph, the inverse was observed; that is, when schema accommodation occurred, increased activity appeared in this network. Taken together, these results indicate that there seems to be a necessary and sufficient relationship between the occurrence of schema accommodation and the increase in activity in the executive network. Therefore, at least in the present experimental conditions, it is suggested not only that deliberation carried out by the executive network can induce schema accommodation but also that schema accommodation would not be possible without it.

By contrast, in both groups, our whole-brain analysis did not result in any clusters that showed significantly smaller signals in the interpretation-changed trials than in the interpretation-unchanged trials. However, considering that the involvement of the vmPFC has been established in schema assimilation, the conceptual counterpart of schema accommodation, it is natural to expect that signals in this area would decrease in interpretation-changed trials (see Introduction). Therefore, we conducted a region of interest (ROI) analysis using a mask covering the medial PFC. In the accommodation group, we observed a cluster in the vmPFC, whose signal was significantly smaller in the interpretation-changed trials than in the interpretation-unchanged trials (Fig. 5 and Table S1). In contrast, in the no-accommodation group, no regions showing decreased activity were found̶whether in the ROI analysis using the medial PFC mask, in the ROI analysis using the clusters identified in the accommodation-group analysis above as a mask, or in the whole-brain analysis.

**Fig. 5.**
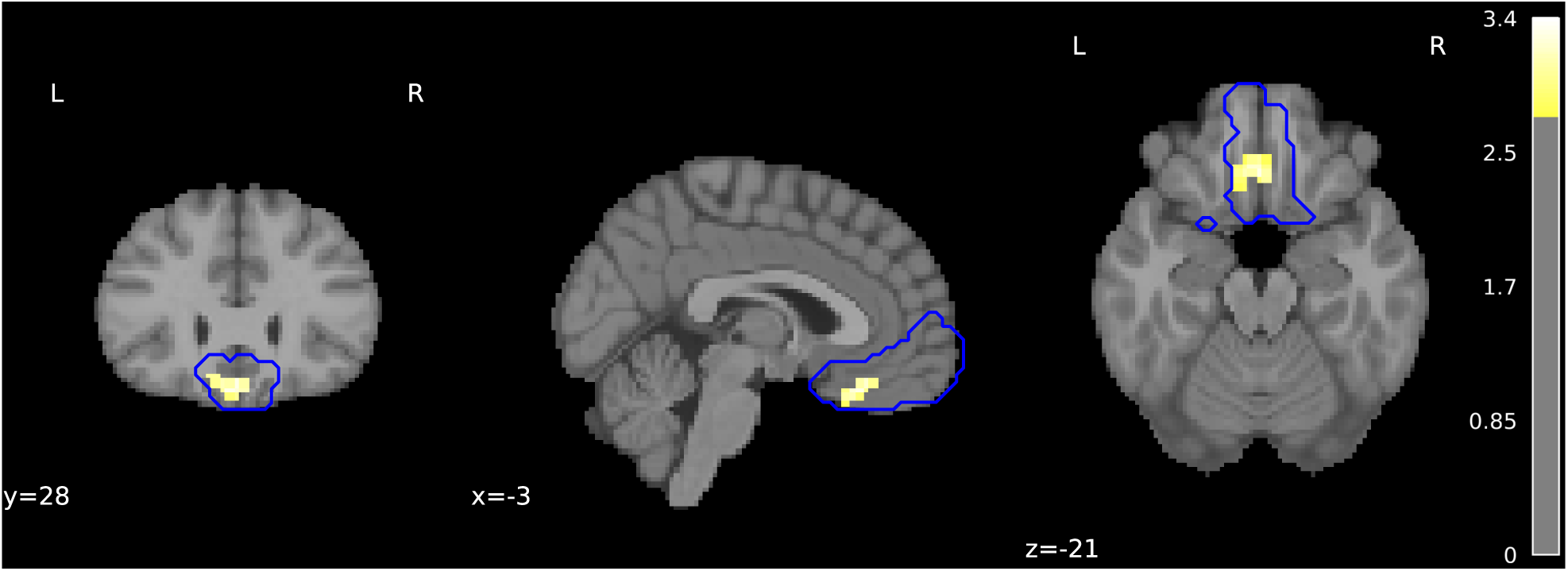
Activity in the ventromedial prefrontal cortex is suppressed during schema reorganization in the accommodation group. Brain regions showing significantly lower activity during interpretation-changed trials compared to interpretation-unchanged trials in the accommodation group are displayed (*p* < 0.05, family-wise error corrected). This result was obtained through a region-of-interest analysis using a mask encompassing the medial prefrontal cortex, indicated by the blue contour line. Voxel-wise t-values are represented using a color-coded scale.

To further investigate the differences in brain activity between interpretation-changed and interpretation-unchanged trials in the accommodation group, we examined psychophysiological interactions using the four clusters found in the whole-brain analysis described above as the seed clusters (Fig. 6 and Table S1). Namely, the blood oxygen level-dependent (BOLD) signal in the seed was set as a physio variable, and the “interpretation-changed vs. unchanged trials” condition was set as a psycho variable. We then explored the areas where the fMRI signal could be explained by the interaction between them. We found clusters in the left and right lateral occipital cortices showing significantly higher connectivity with the seed in interpretation-changed trials (whereas no regions showed higher connectivity in interpretation-unchanged trials). This area has been shown to cooperate with the executive network in tasks such as mental arithmetic and working memory (*43–45*). In particular, this area is considered to play the role of a so-called visual scratch pad (*46*), which is required for deliberate processing. This also suggests that deliberation implemented in the executive network induces the schema accommodation observed in this study.

**Fig. 6.**
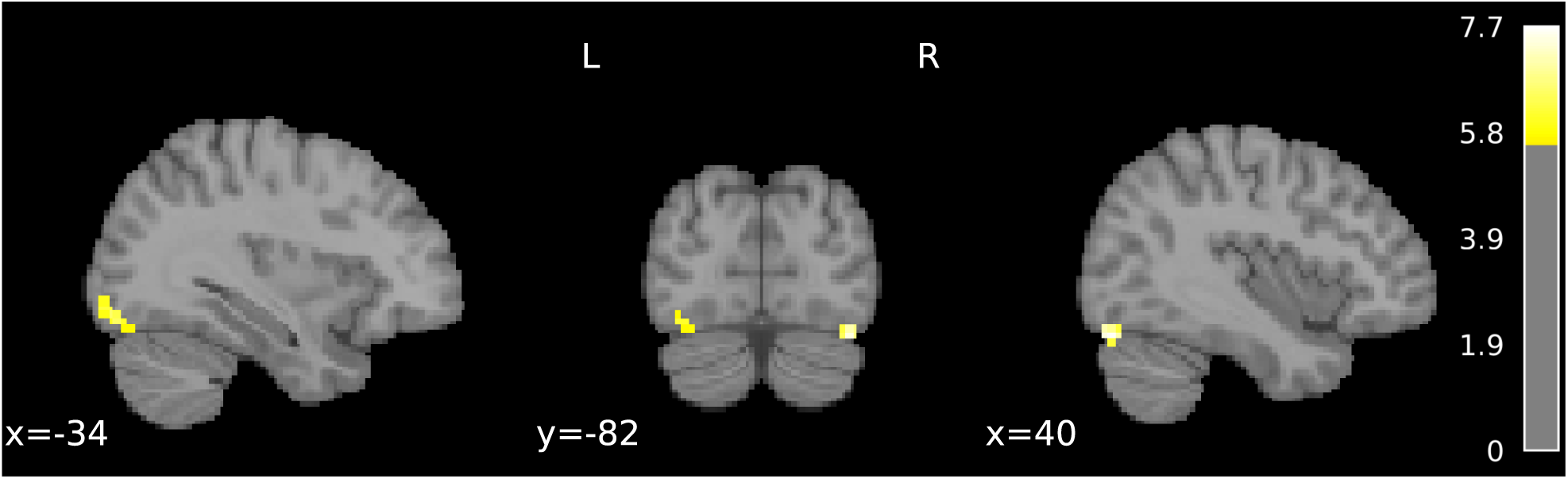
The executive network strengthens functional connectivity with the lateral occipital cortex during schema reorganization in the accommodation group. We examined psycho-physiological interactions using a seed region composed of four clusters—primarily located within the executive network—identified through whole-brain analysis. The figure displays regions showing significantly stronger functional connectivity to the seed during interpretation-changed trials compared to interpretation-unchanged trials within the accommodation group (*p* < 0.05, family-wise error corrected). Voxel-wise t-values are represented using a color-coded scale.

### Maintenance of widespread amplified brain activity might enable schema accommodation

In the previous section, we investigated the differences in brain activity between the interpretation-changed and interpretation-unchanged trials. This was observed in the accommodation group but not in the no-accommodation group. Considering the activity in the interpretation-unchanged trials as a baseline, we assumed that some additional psychological and/or computational factors were reflected in this difference in brain activity. However, in the behavioral analysis described above, we found that the decision durations in the interpretation-changed trials of the accommodation group were shorter than those of the other three cases. Therefore, the participants in this case may have cut more corners than those in the other cases. In other words, activity in the interpretation-changed trials might be considered the baseline, and some psychological and computational factors subtracted from it may explain the differences observed in the accommodation group.

Therefore, we compared the beta estimates, which are assumed to indicate the magnitudes of the brain responses under each condition, for the accommodation and no-accommodation groups (Fig. 7 and Table S2). Specifically, the beta estimates are the coefficients of the explanatory variables estimated by the GLM analysis and reflect the response amplitudes of the BOLD signals. Here, we examined the beta estimates separately for the variables specifying the interpretation-changed trials and those specifying the interpretation-unchanged trials. In both trials, the voxel clusters showed significantly larger beta estimates in the accommodation group than in the no-accommodation group. They are distributed across a wide area of the brain. In contrast, no region showed significantly larger beta estimates in the no-accommodation group than in the accommodation group. These results suggest that regardless of the type of trial, the accommodation group processed information at a greater intensity than the no-accommodation group. Therefore, it is unlikely that the accommodation group in the interpretation-unchanged trials was cutting corners, at least when compared to the no-accommodation group. Furthermore, the overall higher activity of the accommodation group compared to the no-accommodation group may explain the behavioral differences between these groups. In other words, it is possible that the maintenance of this widespread amplified brain activity enables schema accommodation through deliberate processes.

**Fig. 7.**
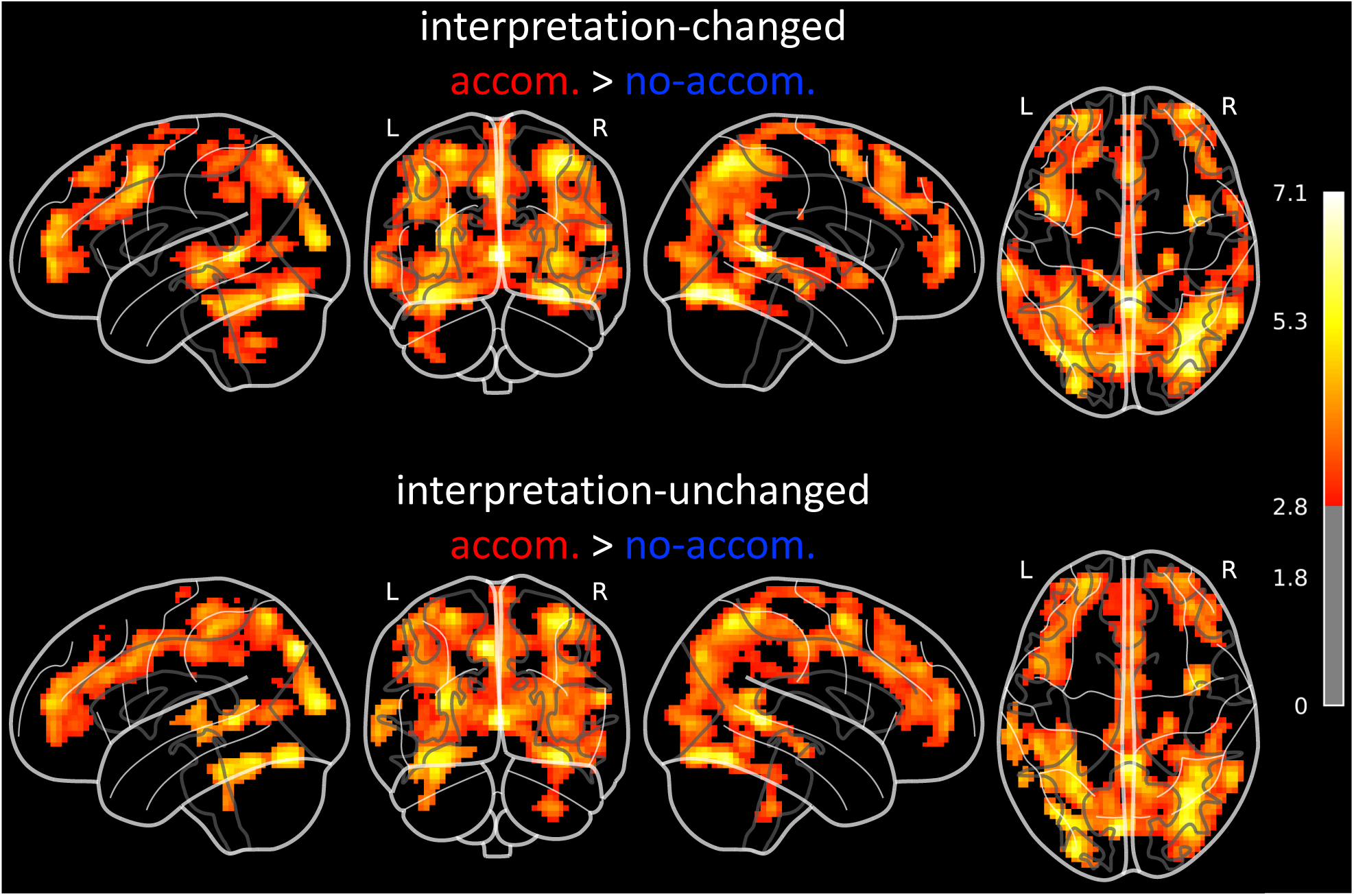
BOLD signal differences between accommodation and no-accommodation groups across trial types. The accommodation group exhibited higher BOLD responses than the no-accommodation group in both interpretation-changed and interpretation-unchanged trials, with these differences widespread across the brain. The maps show the coefficients of the explanatory variables—commonly referred to as beta estimates— estimated for each trial type using whole-brain general linear model analysis. These values were compared between groups. The upper and lower panels display regions with significantly higher beta estimates in the accommodation group compared to the no-accommodation group for interpretation-changed and interpretation-unchanged trials, respectively (*p* < 0.05, family-wise error corrected). Voxel-wise t-values are represented using a color-coded scale.

### Involvement of the caudate nucleus found under a hypothesis on the computational mechanism of schema accommodation

First, let us consider the computational mechanisms underlying the induction of schema accommodation through deliberation, a phenomenon suggested in the above sections. Generally, deliberation is accompanied by a strong sense of agency and requires control processing, such as the manipulation of imagery and symbols in the mind. Furthermore, analyses using network control theory have shown that the executive network, which is responsible for deliberation, is located in brain regions with particularly high modal controllability (*36*), indicating its suitability for difficult control tasks. Those suggest that deliberation is actually “control” in the sense of the control theory (see the supplementary materials (*36*) for details on the control theory). Furthermore, considering the possibility of implementation in the brain, a control based on reinforcement learning (see Ref. (*47*) for details on reinforcement learning) appears to be appropriate because it does not require a clear predefinition of the goal states. When our task is considered from the perspective of control and reinforcement learning, it is natural to assume that the executive network as a controller manipulates the schema structure as a “state.” Thus, from the viewpoint of functional correspondence, the so-called actor-critic reinforcement learning in which the executive network corresponds to the “actor” is assumed. In addition, considering previous findings suggesting that the basal ganglia are the neural substrates of the value function (i.e., the “critic”) of reinforcement learning (*47*, *48*), together with the neuroanatomical evidence of subregional connections between the cortex and basal ganglia (*49*, *50*), we expect the caudate nucleus to be the “critic.” This is because the executive network is primarily interconnected with this subregion of the basal ganglia. See the Discussion section for additional explanation of this hypothesis.

Based on this hypothetical consideration and our results suggesting the involvement of deliberation implemented in the executive network in schema accommodation, it was expected that the accommodation group would show higher activity in the caudate nucleus in interpretation-changed trials. To test this prediction, a GLM analysis was performed using ROI masks on the left caudate nucleus (Fig. 8 and Table S3). As expected, in the accommodation group, voxel clusters were found in the region that showed significantly higher signals in interpretation-changed trials than in interpretation-unchanged trials. No significant clusters were found under the other conditions. This supports the validity of the above hypothesis, claiming that deliberation that induces schema accommodation could be a control process based on actor-critic reinforcement learning.

**Fig. 8.**
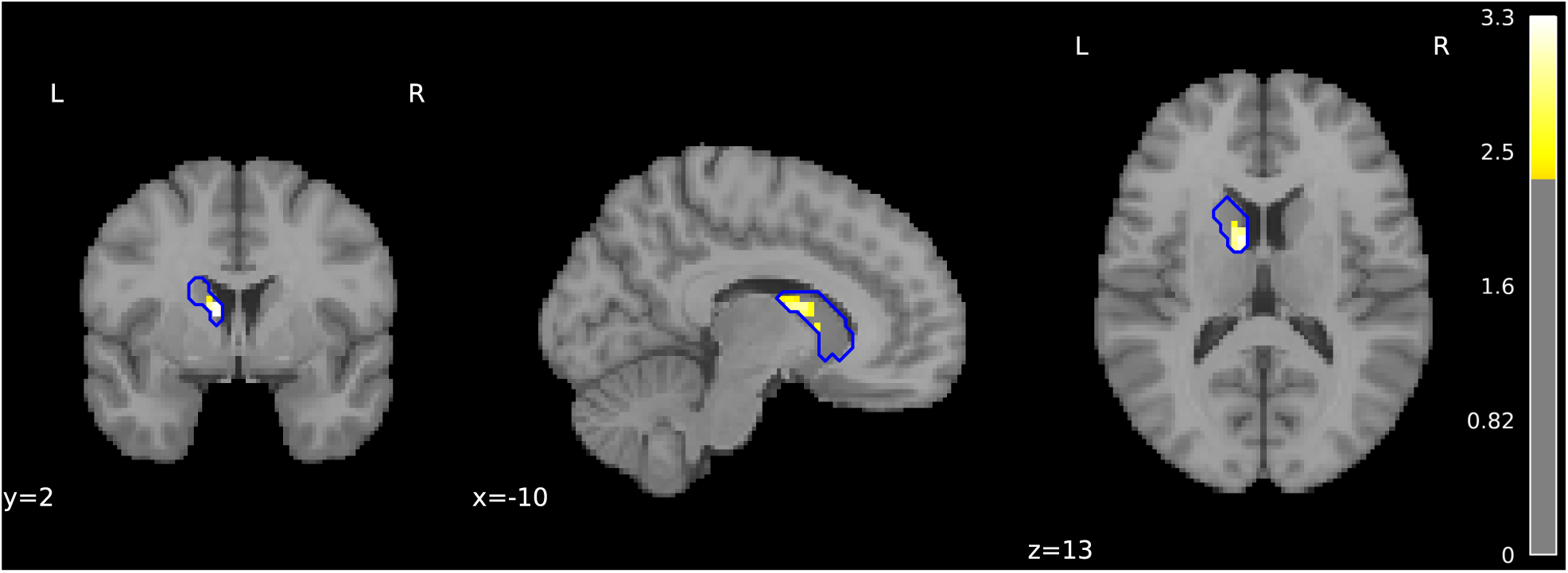
The involvement of the caudate nucleus in the reorganization of schema in the accommodation group. This figure presents the results of a region-of-interest (ROI) analysis using a mask encompassing the left caudate nucleus. The analysis was motivated by a computational hypothesis and anatomical evidence of connectivity between the basal ganglia and the neocortex. The hypothesis posits that reinforcement learning-based control serves as a mechanism of deliberation that induces schema reorganization (see main text for details). The figure highlights the region showing significantly stronger activity during interpretation-changed trials compared to interpretation-unchanged trials in the accommodation group (*p* < 0.05, family-wise error corrected). Voxel-wise t-values are represented using a color-coded scale. The blue contour line indicates the ROI mask.

## Discussion

In the present study, we investigated the neural basis for inducing and/or supporting schema accommodation (namely, the global reorganization of memory structure) and examined its psychological and computational mechanisms. In our analysis, we used the operational definition of schema accommodation. Nevertheless, this seems to have been sufficient to examine the mechanism, as explained below. Participants whose answers to the two-choice questions were reversed before and after the fMRI scan were considered to have undergone schema accommodation (accommodation group), whereas those whose answers were not reversed were considered to have not undergone schema accommodation (no-accommodation group). The behavioral analysis showed that the differences between the groups could not be ascribed to local changes in the schema (i.e., changes in interpretation only at the item level), suggesting that there were some global structural changes underlying the differences. Furthermore, regardless of the trial type, the accommodation group showed increased activity over a wide range of brain regions compared with the no-accommodation group. Therefore, the former group seemed to solve a problem that required a higher cognitive load than the latter. Moreover, only in the former group did fMRI signals and functional connectivity in brain regions associated with cognitive control and attention (FP and CO networks and lateral occipital cortex) (*37*, *38*, *44*, *45*) increase during trials in which changes were made to the schema. Accordingly, taken together, the operational definition of the occurrence of schema accommodation used in this study presumably captures an essential aspect of the phenomenon, namely the global changes in the structure of the schema through mental manipulation.

### Suggested neural substrates playing roles for schema accommodation and assimilation

Using whole-brain analysis of the accommodation group, we identified four voxel clusters that showed stronger activity in interpretation-changed trials. We relate each cluster to previous findings and existing atlases and consider the implications of this observation. As a summary figure of the following discussion, we provide a schematic showing the contrasting neural mechanisms of schema accommodation and assimilation suggested by the findings of this study and previous studies (Fig. 9A).

**Fig. 9.**
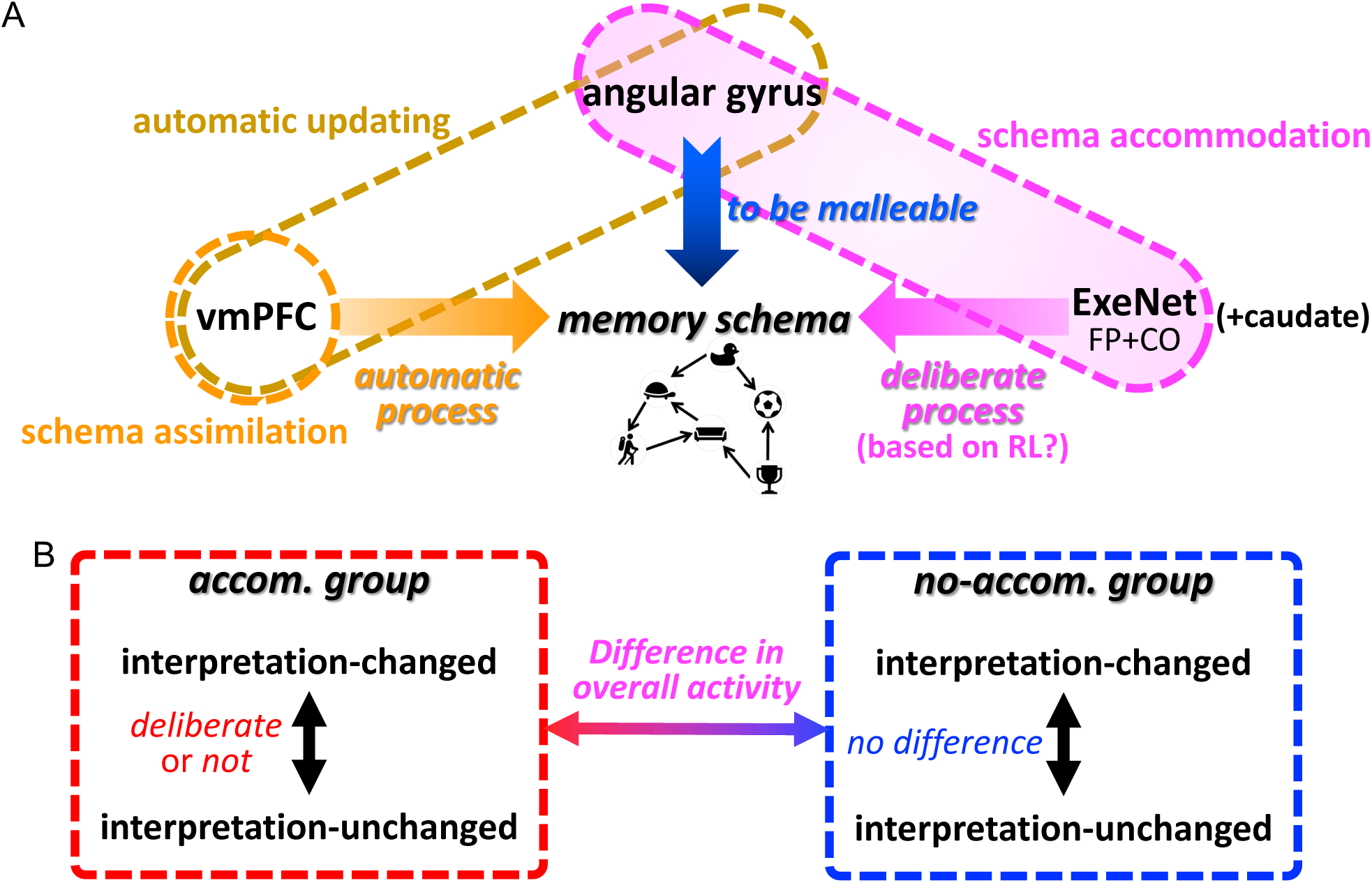
Suggested neural mechanisms of schema updating and summary of the present findings. **(A**) Based on a comparison of prior and current studies, we propose a neural model for schema updating that involves two distinct processes: an automatic process and a deliberative process. The automatic process primarily engages the ventromedial prefrontal cortex (vmPFC), which plays a central role in schema assimilation. The deliberative process, grounded in the executive network, gives rise to schema accommodation together with the schema malleability afforded by the angular gyrus; the executive network underlies deliberation, while the angular gyrus provides the flexibility required for accommodation. This process may be characterized at the computational level as reinforcement learning, with the caudate nucleus, together with the executive network, potentially constituting a key part of its neural implementation. Notably, the angular gyrus also contributes to automatic schema updating through its interaction with the vmPFC. **(B)** Summary of the current study’s findings: In the accommodation group, changes in item-level interpretation within the schema—likely accompanied by reorganizations in the schema’s global structure—were driven by deliberative processing. This was supported by widespread and amplified neural activity, resulting in distinct behavioral and neural differences between the groups.

### Deliberation and the executive network

The largest cluster is observed in the left MFG. Comparing this with the atlas proposed by Yeo et al., which is based on resting-state functional connectivity (*35*), we found that the cluster has substantial overlap not only with Yeo’s FP network cluster but also with Yeo’s default mode network (DMN) cluster that extends into the outer PFC, and the left IPL cluster that we found also spanned both Yeo’s FP network and DMN clusters. Therefore, one may conclude that schema accommodation is carried out through processing, which primarily involves the DMN. However, our analysis did not show any involvement of the precuneus, the core region of the DMN (*51*), and the medial prefrontal cortex (mPFC), a central region of the DMN (*52*), showed rather reduced activity. Moreover, the Yeo atlas is based on resting-state fMRI and does not necessarily capture the properties of task-driven fMRI responses. Additionally, because it identifies clusters by hard clustering, which does not allow the sharing of elements, cortical gradients that have recently attracted attention have been ignored (see Ref. (*53*)). In fact, an analysis of cortical gradients based on fMRI responses to a set of parameterized cognitive control tasks suggests that regions contributing to higher-order cognitive control include, in particular, those classified into the DMN in the Yeo atlas (*54*). In addition, gradients corresponding to the contribution to cognitive control have been found in both the PFC and PPC (*54*), indicating that the contribution to the function of the region is not all-or-nothing-like. Given these considerations and the details of our experimental task requiring effortful mental manipulation, it is reasonable to assume that information processing, deeply related to executive control, underlies the observed phenomena. However, there are reports of cooperation between the DMN and executive network in some tasks (*55*) and decomposition of the DMN in schema-related tasks (*56*). Both networks may have collaborated in this study.

Significant clusters were also found in the left medial and lateral SFGs. In general, the medial SFG is part of the CO network (*34*). However, the medial SFG cluster that we observed was mostly included in the FP network cluster in the Yeo atlas. Nevertheless, both the CO and FP networks comprise the executive network and are functionally closely associated with each other (*37*, *57*, *58*). Therefore, whether it is part of the CO or the FP network does not affect our consideration of the psychological and/or computational mechanisms of schema accommodation. Meanwhile, the left lateral SFG cluster that we observed overlapped with the Brodmann area 8. Because this cluster was lateralized to the left hemisphere, it was considered to cooperate with the other observed clusters that comprise the executive network, which were also lateralized to the left. This is consistent with several studies suggesting the involvement of this area in executive functions (e.g., Ref. (*59*)).

Many previous studies have shown that the executive network engages in higher-order intellectual and rational information processing (*40–42*). As such processing is generally accompanied by intentionality, it is appropriate to interpret it as deliberate processing. This is consistent with the other findings and task settings in the present study. Notably, the observed increase in functional connectivity between the executive network and the lateral occipital cortex adds further mechanistic details to this interpretation. This area is known to co-activate with regions involved in executive functions during working memory tasks (*44*, *45*). In particular, it acts as a visuospatial sketchpad (*46*). Schema accommodation requires the manipulation of one’s schema, and the visuospatial sketchpad can be used as a place to do this through trial and error (see Ref. (*60*)). This further supports the interpretation that the increased activity in the executive network observed in this study reflects deliberation. In light of these considerations, we assume that the extent of activity in the executive network is a marker of the degree of deliberation. Schema accommodation occurred in the group whose magnitudes of marker activity were greater in the interpretation-changed trials than in the interpretation-unchanged trials. In contrast, this did not occur in the group in which the magnitudes of the deliberation markers did not differ between trial types. Therefore, we propose that schema accommodation is induced by a deliberate process implemented in the executive network (Fig. 9A, right; Fig. 9B, left).

### The angular gyrus might make schema editable

As mentioned previously, the left IPL cluster found in this study spanned both the FP and DMN network clusters of the Yeo atlas. The latter anatomically corresponds to the angular gyrus. The involvement of the angular gyrus is important for updating memory schema. Several neurophysiological experiments using tasks in which participants were presented with naturalistic videos or spoken audio and asked to understand the story (i.e., to construct a new schema) and then update their understanding through exposure to twists, spoilers, or hidden connections have consistently reported the involvement of the angular gyrus (*26*– *29*). In particular, an intervention experiment using transcranial magnetic stimulation showed that the angular gyrus played a causal role (*29*). Unlike in the present study, the schema changes examined in these studies were not cognitively demanding. In fact, no resistance to changes in the schema was observed (i.e., it was not schema accommodation). The tasks of the present and previous studies have in common the fact that the complex and rich pre-existing schemata must be changed to a substantial extent. Therefore, to update memory schema, the angular gyrus may play a role in making it malleable and editable (Fig. 9A, center).

### The vmPFC might be responsible for an automatic aspect of memory dynamics

In the present study, ROI analysis using the mPFC mask showed that vmPFC activity decreased in the interpretation-changed trials in the accommodation group. On the other hand, it has been almost established that increased activity in this region plays a role in schema assimilation (*3*, *4*, *9*) that is often viewed psychologically in contrast to schema accommodation. Thus, schema accommodation and assimilation also contrast in terms of the brain activity involved. Interestingly, the experiments mentioned above, which used a task to update understanding through twists, spoilers, or hidden connections, also reported increased activity in this region (*26*, *27*). The key difference between the experiments in these previous studies and ours was that the participants in the former study were not subjected to cognitive difficulties in performing these tasks. Instead, they are easily performed as automatic unconscious processes. Therefore, the vmPFC appears to be involved in automatic aspects of memory dynamics (see also Ref. (*22*)) (Fig. 9A, left). Thus, the decreased activity in the vmPFC observed in the present study may reflect the suppression of autonomous memory processing in this area, which may be associated with FP and CO network activity (*61*). From a computational perspective, in both the previous and present experiments, the participants transitioned their schemata from the initial state to the goal state. However, they presumably did so along a routine or guided path in the former case, but by trial and error in the latter. As discussed below, the latter can be processed based on reinforcement learning. Overall, a contrasting picture of unconscious and automated schema assimilation versus strongly conscious and deliberate schema accommodation was presented (Fig. 9A).

### Functions of widespread amplified brain activity in schema accommodation and its physiological mechanisms

The most striking difference between the accommodation and no-accommodation groups was the magnitude of overall brain activity, regardless of the trial type. The former group showed greater widespread brain activity, suggesting that they were internally solving more cognitively demanding problems or that the cognitive efforts of the latter group were insufficient. From a mechanistic standpoint, this widespread increase in brain activity could support deliberate processing with sufficient effort by the executive network (Fig. 9B).

Because few previous studies have associated such an overall increase in brain activity with the realization of specific task execution, further interpretation of this phenomenon requires the accumulation of future research findings using similar analyses. In particular, it is important to investigate whether this phenomenon reflects a transient psychological and/or neural state or whether it is due to an individual’s long-term intrinsic properties. In addition, it is important to theoretically evaluate the effects of this overall amplification of activity. The bifurcation theory in dynamical systems may provide valuable insights. This suggests that, depending on the macroscopic parameters, the system’s possible states can change qualitatively. For instance, it has been demonstrated that solutions exhibiting stationary waves (i.e., continuous attractors) emerge in a neural field model, depending on the intensity of random background activity (*62*). Such qualitative changes may account for the differences observed between the groups in the present study.

### Reinforcement learning-based control as a possible computational mechanism of deliberation inducing schema reorganization

We hypothesized that the computational mechanism for schema accommodation investigated in this study involves controlling schema structure in a control-theoretic sense. This is based on an analysis of the psychological concept of deliberation and previous research showing that the executive network is well-positioned to perform difficult control tasks, especially those that require states to cross high potential barriers (*36*). In addition, because it is difficult for the brain to know in advance the goal structure of a schema that can accept novel information incongruent with the existing one, we considered reinforcement learning-based control to be appropriate because it does not require such prerequisite knowledge (see Ref. (*47*) for details on reinforcement learning). Multivoxel pattern analysis of fMRI data and diffusion-weighted MRI studies of structural plasticity suggest that the schema is not contained within the executive network (which is a possible controller), but is in regions such as the mPFC, precuneus, and angular gyrus (*63–65*). From the viewpoint of reinforcement learning, the functional and/or structural states of these regions can be regarded as “states” to be controlled. Furthermore, studies on the neural basis of reinforcement learning suggest that the value function is implemented in the basal ganglia (*47*, *48*). Subregions of the cortex and basal ganglia have specific anatomical connections; in particular, the region corresponding to the executive network is known to have connections with the caudate nucleus (*49*, *50*). Therefore, in the present case, the value function was considered to be located in the caudate nucleus. Thus, actor-critic reinforcement learning is assumed as the computational mechanism of deliberation as control, in which the executive network is the “actor” (i.e., the controller of the state), the caudate nucleus is the “critic” (i.e., the state value function), and the schema structure existing in the aforementioned regions is the “state.” Our analysis, based on this assumption, showed the involvement of the caudate nucleus.

In neuroimaging experiments using typical reinforcement learning tasks, it has been reported that the state value function is not located in the dorsal striatum (including the caudate nucleus) (*66*, *67*), but rather in the ventral striatum (*68*). This discrepancy is presumably due to differences in the anatomical connections between these subregions of the striatum and the regions involved in the task. However, other possibilities cannot be ruled out. Further studies are needed to confirm the theory that deliberation is a control process based on actor-critic reinforcement learning implemented in the abovementioned neural bases.

Given the lack of a consensus on the computational mechanism of deliberation, one can understand the additional significance of proposing this theory of memory reorganization. In other words, this theory contributes to elucidating the computational mechanisms of not only memory reorganization but also deliberation itself. It should be mutually beneficial such that the elucidation of deliberation may reveal an unknown mechanism of memory reorganization, and that the subject of memory reorganization may provide an appropriate experimental system as a representative model of deliberation to examine its computational theory.

### Long-term memory-related processing in the lateral prefrontal cortex

The lateral PFC (lPFC), which is part of the executive network, has been studied mainly in the context of short-term memory (or working memory) (see Ref. (*69*)). In contrast, studies examining its role in long-term memory have been scarce. Nevertheless, there are several important findings related to this issue. In particular, evidence suggests that the lPFC contributes to encoding information into long-term memory through actively manipulating it in the mind during the maintenance period (*70*).

Additionally, several studies have suggested the involvement of the lPFC in modifying established schemata, such as those examined in the present study (*22*, *23*, *71*, *72*) (see also Ref. (*21*)). In one of these studies, the participants engaged in an association task between facial images and the names of real cities (*23*). The results showed that updating involved an increase in lPFC activity. We interpret this in terms of the executive control functions of the lPFC. There are two important points to note regarding this study. First, as in the present study, resistance to updating the association was observed during the task. Second, because the city names used in the experiment exist and are well-known, the associations that they formed might have been rich and complex schemata in which this information was incorporated. If so, participants were required to perform high-level manipulations of the schema structures to update the associations. In other words, it is possible that, in this experiment, they engaged in schema accommodation using executive functions, as was the case in the present study. This may explain the involvement of the lPFC. In addition, it has been shown that lPFC activity is more strongly involved in changing social norms, which are more resistant to persuasion, than in changing mere beliefs, which are less resistant (*72*). Furthermore, this study also showed increased IPL activity in the social norm change condition, which was shown to be involved in schema accommodation in the present study.

A theoretical framework explaining the findings of these studies, including ours, is required to clarify the role of the lPFC in memory acquisition and updating.

### Implications of the findings to dynamical and systemic aspects of memory schema

The primary motivation for this study was to understand the self-organization principle of schema structures, namely, the law governing the temporal evolution of memory schemata through online and offline experiences (e.g., learning and thoughts). Since the schema shapes perceptions, thoughts, and actions, the law binds the capabilities of each individual and all humanity; that is, what kind of order and integrity of cognition, knowledge, and intelligence can we possess?

The operation of deliberation on memory reorganization suggested in this study has fundamentally distinct implications for the law governing the temporal evolution of memory from those in previous studies. In most neuroscientific, and psychological experiments, participants are prevented from deliberating. For instance, experimenters loaded participants’ working memory to prevent deliberation (for example Ref. (*73*)). This is because of the current status of neuroscience, where deliberation has not yet been well defined in a computational sense, and our everyday intuition that we can do a variety of things through deliberation. That is, if there is strong deliberation during task performance, its versatility or universality can lead to arbitrary results, and thus, to a failure to identify the law of the phenomenon that we wanted to examine.

The central role of deliberation in schema accommodation suggested in this study has important and difficult consequences for understanding the laws governing the dynamics of the schema and the intelligence created through it. This means that, to understand them, we must deal with deliberation. Therefore, our suggestion that the induction of schema accommodation through deliberation is based on reinforcement learning is particularly important because it provides a promising theoretical and experimental guide for elucidating deliberation.

### Limitations and future directions

In the present analyses, we found a necessary and sufficient relationship between the occurrence of schema accommodation and greater activity in the executive network during interpretation-changed trials. However, this was observed only under the conditions examined in this study. It is plausible that executive network activity can induce some types of schema accommodation; however, whether it is generally required for schema accommodation to occur remains unclear. Both schema assimilation and accommodation may be located on a continuum, with the amount of change in the schema structure as the axis. Therefore, schema assimilation may cause a small but non-negligible reorganization of the schema structure. In such cases, repeated schema assimilation, which does not require executive network activity, can result in global changes in schema structure, similar to schema accommodation. In addition, while we used an emotion-neutral task in this study, it is presumably common for people’s worldviews (of which schemata are their components) to be changed by events that strongly appeal to their emotions. Therefore, schema accommodation in emotionally arousing situations may involve mechanisms that differ from those observed in this study.

Considering the results of the behavioral and fMRI analyses and the properties of the task together, we concluded that the accommodation group underwent some global changes in their schema structures. Studies on representational similarity analysis, graph-theoretic analysis of free associations of words, object similarity evaluation, and machine learning-based prediction of object memorability imply that we have schemata featuring characteristic global structures (*6*, *74–76*). It has also been shown that their structures are involved in recognition, retrieval, creativity, etc. (*5–8*). These studies highlight the need for a new research discipline that focuses on the global structures of memory schemata. This study, which approaches the principle of self-organization of structures, can provide a basis for this discipline. However, our study did not directly observe the schema structures constructed in the individuals or their changes. Therefore, future work is needed in which the task is revised so that behavioral outcomes that more directly reflect each participant’s schema are observable, and behavioral and fMRI experiments are conducted using the revised task.

Another limitation of this study is that the relationship between deliberation and executive network activities is unclear. This is not a trivial issue; rather, it is a fundamental challenge. However, as discussed in the previous section describing the significance of this work, the experimental systems and computational hypotheses developed here already provide a concrete route toward resolving this problem.

Beyond addressing such unresolved issues, these experimental paradigms and hypotheses, together with the present results, should provide the basis for a newly developing research field investigating the global structure of memory. It is hoped that this will lead to the elucidation of memory as a self-evolving complex system̶flexible, adaptive, and creative, like a living organism, yet sometimes more accurate than artificial machines, and has diversity among individuals, yet systematicity and integrity in each to their own way.

## Materials and Methods

### Participants

Thirty-four subjects (14 females and 20 males; mean age 20.6 years; age range, 18–24 years) with no history of neurological or psychiatric diseases participated in this study. All the participants were native Japanese speakers with normal or corrected-to-normal vision. This study was conducted in accordance with the recommendations of the institutional ethics committees of Tokai University and Tamagawa University. Written informed consent was obtained from all subjects. All participants provided written informed consent in accordance with the Declaration of Helsinki. The protocol was approved by the institutional ethics committees of Tokai University and Tamagawa University.

### Experimental procedures

To investigate which neural mechanisms induce and/or support schema accommodation, it is first necessary for participants to acquire schemata that are new to them and are as rich as everyday memories or naturalistic knowledge. This is because, as noted above, schema accommodation is considered a phenomenon in which the structure changes; therefore, it is necessary to study schemata with sufficient structural complexity, but this must be under the experimenter’s control. In addition, as mentioned above, schemata that individuals naturally possess are resistant to accommodation; therefore, the schema to be acquired in this experiment must also be firmly established enough to have such resistance. However, at the same time, schema accommodation must occur with sufficient efficacy against its resistance to enable it to be sufficiently detected in the measurement conditions to elucidate its neural basis using fMRI. To this end, we conducted an fMRI experiment using a newly developed experimental paradigm called the reversal description task (Fig. 1B). Here, we describe the procedure with an in-depth explanation of the reasons behind each step.

The experiment was conducted over two consecutive days. On the first day, only the behavioral experiment was conducted. Participants were given a document and asked to read it carefully while taking notes and were then required to write a text summarizing their understanding. The document was presented on an A4 paper sheet printed on both sides. The participants were also given another sheet of A4 paper as a notepad on which they wrote their notes using a ballpoint pen. Writing was performed in a Microsoft Word file using a typical PC display, keyboard, and mouse. During this writing session, they were not allowed to see the document but were only allowed to refer to their notes. However, during the reading session, we did not prohibit them from writing anything on the document paper. This set of reading and writing was repeated twice. The first reading and writing session lasted 15 minutes each (30 minutes in total) and the second reading and writing session lasted 10 minutes each (20 minutes in total). As I will explain in more detail later, the document was very esoteric but possible to comprehend in principle and was insightful (in other words, highly pedantic), and therefore, to read it and then write their understanding required actively creating an interpretation in their minds. This means that, by setting up a situation that demanded active interpretation, each participant was forced to construct a new schema in his/her mind that was sufficiently rich and robust.

At the beginning of the second day, this set of reading and writing sessions was repeated, each lasting 10 min. After that, participants were asked to answer a two-choice question (“A” vs. “B”) in which their answers depended on their interpretations of the document (i.e., on the schemata constructed in their minds) (For details of the question, refer to the following section). Then, the participant who answered “A” was told that “B” was the correct answer, and the participant who answered “B” was told that “A” was the correct answer. Furthermore, we told each participant that after the MRI measurement, he/she would again write a text summarizing his/her understanding considering that the opposite answer was the “correct,” and that to do this, during the fMRI session, he/she would have to change the interpretation so that the “correct” answer would be his/her own answer. In addition, they were informed that they would be given a task to facilitate this change of interpretation during the session. Thus, since the procedure of this experiment requires the participants to reverse their original interpretation and then describe a text based on the reversed interpretation, we named it the “reversal description task.”

During the fMRI session, 40 sentences extracted from the document were presented sequentially in randomized order among the participants (Fig. 1C). Additionally, the “correct” answer was displayed at the top of the screen as a reminder. Participants read each sentence, rethought it, and reported whether their interpretation of that sentence had changed. We asked them to report whether the interpretation of the sentence presented in front of them had changed, not the interpretation of the entire document, the answer to the two-choice question mentioned above, or the interpretation of a sentence other than the one presented at that moment. Thus, the question concerns an item-level change in the schema. Specifically, participants internally recognized whether their interpretation of the sentence had “changed” or “not changed,” and pressed the button at the moment when the recognition was reached. Therefore, this part was self-paced. The buttons consisted of “Up,” “Down,” “Left,” and “Right,” but only the “Up” button was used here, regardless of which was recognized. In the next scene, the recognition outcome was reported by moving the underline among “changed,” “unchanged,” and “unknown/other” by pressing the “Up” button, and by submitting it by pressing the “Right” button, again in a self-paced manner. Subsequently, the part in which the sentence was viewed passively continued for six seconds. Then, a blank part of 2 s followed, and the next trial was moved to. For the trials in which “changed” was selected, some changes in the schema were considered to have occurred, and for the trials in which “unchanged” was selected, no changes in it were considered to have occurred. Therefore, the main interest of this study was to compare the brain activity occurring in these two types of trials.

After the MRI session, the participants again wrote about their current understanding of the document. This task lasted for 10 min. They were no longer able to read their notes, but some words that appeared in the document were presented to help them recall the content. Subsequently, participants answered the same two-choice questions as before. They were instructed by the experimenter to answer honestly.

### The document and the two-choice question

As mentioned above, in this experiment, participants constructed a new schema by reading an esoteric document that required active interpretation. The document was carefully selected to meet the following criteria: First, it had to be difficult yet, in principle, comprehensible. Accordingly, we targeted a real-world document that has been read and debated by many readers. However, it had to be a novel subject that participants (mostly university students) had never studied or thought about before. On the other hand, if the subject matter was so remote that it was unrecognizable to them, it was not appropriate; some degree of familiarity with it needed to be expected. The document also had to be logically challenging; however, the words comprising it had to fall within the vocabulary of typical university students. In light of these considerations, we chose the theories of Niklas Luhmann, a theoretical sociologist known for his esoteric, pedantic, but thought-provoking writings, as our subject matter. See Ref. (*77*), which is an English translation of Luhmann’s main work. Because the document used in the experiment had to be self-contained, clippings from Luhmann’s writings were inappropriate. Therefore, we used the section on “Complexity” in the Japanese translation of the glossary “GLU: Glossar zu Niklas Luhmanns Theorie sozialer Systeme” (*30*) as our target document. This is also esoteric and insightful enough. The document presented to the participants comprised the first 58 of the 80 lines of the entire section, and its length was 1,483 words in Japanese. Note that Japanese text is not separated by spaces between words; therefore, we performed a morphological analysis using the MeCab Python module (mecab-python3 v1.0.10, https://taku910.github.io/mecab/) with the dictionary unidic-lite 1.0.8 (https://github.com/polm/unidic-lite) set to identify the words in advance. Although some of the participants were sociology majors, none had previously studied Luhmann’s theory.

The two-choice questionnaire consisted of a problem statement, options, and confidence rating. They were presented in Japanese. The problem statement was “The level of system complexity changes when the selectivity of the various relationships that are structurally realized changes. Systems can increase their own complexity in relation to the increase in environmental complexity. Why is this?” The options were “A) Because the system responds to the complexity of the environment” and “B) Because the system does not respond to the complexity of the environment.” The confidence level for each choice was self-assessed using the following four-point scale: “1. Just guessing”, “2. Not so sure, but I think so”, “3. I think so”, “4. Surely, I think so”. See Fig. 3C for an evaluation of these choices using deep natural language processing and network theory. Briefly, their semantic centralities were relatively high (though not the highest) compared with those of the sentences presented during the fMRI session. In other words, they were related to the subject matter of the document. Also, note that their degrees of centrality for “A” and “B” were almost the same. Details of the analysis are described later.

### MRI scanning

MRI data were acquired using a 3-Tesla Siemens Prisma Fit MRI scanner (Siemens, Germany) with a 32-channel matrix head coil. Functional scans were obtained with gradient-echo T2*-weighted echo-planar images (EPI; repetition time = 1000 ms; echo time = 26.40 ms; flip angle = 65 degrees; slice thickness = 2.4 mm; the number of slices = 56; acquisition matrix = 80 x 80; voxel size = 2.4 mm x 2.4 mm x 2.5 mm). The slice order was interleaved. Structural scans were obtained with T1-weighted GRAPPA images (repetition time, 2500 ms; echo time = 2.24 ms; flip angle, 8 °; inversion time = 1000ms,; number of slices, 224; voxel size = 0.8 mm x 0.8 mm x 0.8 mm).

### MRI data preprocessing and statistical analyses

Standard preprocessing procedures (slice time correction, realignment, reslicing, coregistration, segmentation, normalization, smoothing (full width at half-maximum (FWHM) = 8 mm)), first-level analysis of functional data, and PPI analysis were performed using SPM12 (The Wellcome Centre for Human Neuroimaging, England, UK). The MRI images of all participants were aligned to the Montreal Neurological Institute (MNI) standard brain. Statistical analyses of the MRI images were performed in the MNI space. Therefore, the coordinates reported in this study are within this space. For higher-level analysis, we used the threshold-free cluster enhancement (TFCE) method (*78*) implemented in FSL 5.0.6 (FMRIB Software Library). SnPM analysis was also performed using the SPM add-in software; however, because the main results were qualitatively similar to those of TFCE, only TFCE was used thereafter.

The explanatory variables for the general linear model in the first-level analysis included binary variables (representing the intervals for rereading and rethinking the presented sentences, intervals for reporting whether the interpretations changed, and intervals for passively viewing the sentences), real-valued variables representing the six motion parameters identified by the motion correction procedure, and the intercept. We used three variables for the rereading-and-rethinking intervals, corresponding to the subsequent reports (“changed” vs. “unchanged” vs. “unknown/other”). Therefore, the design matrix comprises 12 columns. The canonical hemodynamic response was used to deconvolve the fMRI signals; model derivatives were not used. Global normalization was not performed. The masking threshold was set to 0.1. Serial correlations in the fMRI time series were accounted for using the AR(1) model.

The resulting statistic maps of the rereading-and-rethinking intervals corresponding to “changed” and “unchanged” were subjected to subsequent higher-level analyses. In the analyses, paired t-tests were performed using the FSL randomise function, and significant clusters were identified using TFCE. Multiple comparison correction was applied using the “FWE-corrected” method to control for family-wise error rates. The number of permutations was set to 5,000. Variance smoothing was applied with a sigma of 5 mm to increase statistical power.

In addition to the whole-brain analysis, we also performed analyses on predefined regions of interest (ROIs). The ROI mask of medial PFC (mPFC) was created by applying the maximum filtering to the left and right “PFCvm” parcels of the MarsAtlas cortical parcellation (*79*) with a 3-mm radius sphere as a kernel, using the FSL fslmaths function. For the ROI mask of the caudate nucleus, the left “Caudate” segments of the MarsAtlas subcortical structures (*80*) were used.

In addition, to compare the magnitudes of the blood oxygen level-dependent (BOLD) signal responses between the accommodation and no-accommodation groups, we conducted between-group comparisons of beta maps. This was done for each of the maps corresponding to the “changed” and “unchanged” variables, respectively. Analyses were performed using unpaired t-tests with the FSL randomise function. The other conditions were the same as those described above.

Statistical tests other than MRI data were conducted using the Python modules Scipy v1.10.1, Statsmodels v0.14.0, and Pingouin v0.5.4. Semantic analysis of the sentences presented to the participants in the fMRI session was performed using BERTScore (*31*). The BERTScore is a method for quantifying the semantic similarity between sentences using the encoding of sentences by BERT, a Transformer-based deep neural network. This was shown to correlate well with manual evaluations. In this study, we calculated these semantic similarities using a pre-trained model in the Python module bert-score v0.3.13 (https://github.com/Tiiiger/bert_score), linearly normalized them so that their minimum and maximum values were 0 and 1, respectively, and then used them as edge weights to construct an inter-sentence network. The eigenvector centrality of each sentence in the network was calculated using NetworkX v3.2.1. The centralities were divided into interpretation-changed trials and interpretation-unchanged trials for each participant and averaged. The resulting values were used for the statistical tests. Similarly, the BERTScore was also used to calculate the semantic similarities between the sentences presented during the fMRI session and the two sentences in the question (“A” and “B”). That is, for each sentence in the fMRI session, the similarity corresponding to each of “A” and “B” was calculated. However, these two values are almost identical. Therefore, the average value of them was used in the statistical analysis.

## Acknowledgments

We thank T. Kinjo, S. Tsuchida, and M. Osawa for technical assistance. We would like to thank Editage (www.editage.jp) for English language editing.

## Funding

Leading Initiative for the Excellent Young Researchers (MEXT, Japan) (HK)

Grant-in-Aid for Scientific Research (C) (21K07264) (HK).

Joint Research Program implemented at Tamagawa University Brain Science Institute (MEXT, Japan)

## Author contributions

Conceptualization: HK, JK, KM

Methodology: HK, JK, KM

Investigation: HK

Visualization: HK

Supervision: KM

Writing—original draft: HK

Writing—review & editing: HK, JK, KM

## Data and materials availability

All data needed to evaluate the conclusions in the paper are present in the paper and/or the Supplementary Materials. Source data have been available in the OSF repository.

Supplementary Materials

---

**Table S1.** Significant clusters identified in the main analysis.

| Region | x | y | z | number of voxels | p-value (corrected) |
| --- | --- | --- | --- | --- | --- |
| Whole-brain analysis (interpretation-changed > unchanged) |  |  |  |  |  |
| Left middle frontal gyrus | -40.4 | 10.6 | 50.9 | 166 | 0.0068 |
| Left medial superior frontal gyrus | -2.13 | 25.5 | 55.4 | 111 | 0.0254 |
| Left inferior parietal lobule | -42.9 | -59.5 | 51.8 | 38 | 0.0246 |
| Left lateral superior frontal gyrus | -16.7 | 38.2 | 48.3 | 14 | 0.0464 |
| Region-of-interest (ROI) analysis in medial prefrontal cortex (interpretation-changed < unchanged) |  |  |  |  |  |
| Left ventromedial prefrontal cortex | -2.69 | 27.6 | -21.3 | 59 | 0.0302 |
| Psycho-physiological interaction analysis (interpretation-changed > unchanged) |  |  |  |  |  |
| Left lateral occipital cortex | -32.7 | -86.7 | -13.3 | 29 | 0.0362 |
| Right lateral occipital cortex | 39.5 | -78.9 | -21 | 18 | 0.03 |

The x, y, z values indicate the center of gravity of each cluster in the MNI coordinate system. The number of voxels is calculated under the condition of 3 mm isotropic voxels. The p-value (family-wise error-corrected) indicates the minimum value within the cluster.

**Table S2.** Significant clusters identified in beta map analysis.

| Region | x | y | z | number of voxels | p-value (corrected) |
| --- | --- | --- | --- | --- | --- |
| Accom. group > no-accom. group (interpretation-changed > unchanged) |  |  |  |  |  |
| Right precuneus | 4.09 | -56.9 | 14.3 | 4954 | 0.0056 |
| Left middle frontal gyrus | -33.7 | 32.7 | 28.1 | 820 | 0.0198 |
| Right frontal pole (lateral) | 34.2 | 52.4 | 13.8 | 362 | 0.0258 |
| Right paracingulate gyrus | 1.73 | 29.2 | 41.2 | 324 | 0.0286 |
| Left lateral occipital cortex | -27.6 | -88.7 | 17.7 | 101 | 0.029 |
| Right middle frontal gyrus | 38.6 | 5.7 | 56.4 | 82 | 0.0348 |
| Right frontal pole (medial) | 2.21 | 55.6 | 12.4 | 57 | 0.0458 |
| Left parahippocampal gyrus | -17.7 | -30.2 | -22 | 11 | 0.0424 |
| Left cingulate gyrus | 0 | -22 | 27.5 | 2 | 0.05 |
| Accom. group > no-accom. group (interpretation-changed < unchanged) |  |  |  |  |  |
| Right cingulate gyrus | 6.68 | -20.7 | 25.6 | 5447 | 0.011 |
| Left fusiform gyrus | -35.2 | -47.2 | -24.4 | 237 | 0.0252 |
| Left lateral occipital cortex | -27.4 | -88.4 | 18.7 | 158 | 0.0234 |
| Left fusiform gyrus | -31.3 | -74.3 | -16.1 | 152 | 0.0256 |
| Left middle temporal gyrus | -64.2 | -28.1 | 2.01 | 92 | 0.0418 |
| Right middle frontal gyrus | 38.7 | 4.2 | 56.6 | 88 | 0.032 |
| Right angular gyrus | 56.4 | -50 | 16.3 | 72 | 0.0342 |
| Left angular gyrus | -57.7 | -51.2 | 11.5 | 19 | 0.045 |

The x, y, z values indicate the center of gravity of each cluster in the MNI coordinate system. The number of voxels is calculated under the condition of 3 mm isotropic voxels. The p-value (family-wise error-corrected) indicates the minimum value within the cluster.

**Table S3.** Significant clusters shown from analysis based on a computational hypothesis.

| Region | x | y | z | number of voxels | p-value (corrected) |
| --- | --- | --- | --- | --- | --- |
| left caudate region-of-interest analysis (interpretation-changed > unchanged) |  |  |  |  |  |
| Left caudate nucleus | -12 | 0.77 | 15.7 | 21 | 0.0182 |
| Left caudate nucleus | -8.36 | 10.3 | 3.3 | 9 | 0.0356 |

The x, y, z values indicate the center of gravity of each cluster in the MNI coordinate system. The number of voxels is calculated under the condition of 3 mm isotropic voxels. The p-value (Family-wise error-corrected) indicates the minimum value within the cluster.

